# Probing the mechanism of Cbl-b inhibition by a small-molecule inhibitor

**DOI:** 10.1101/2023.05.05.539612

**Authors:** Serah Kimani, Sumera Perveen, Magdalena Szewezyk, Hong Zeng, Aiping Dong, Fengling Li, Pegah Ghiabi, Yanjun Li, Irene Chau, Cheryl Arrowsmith, Dalia Barsyte-Lovejoy, Vijayaratnam Santhakumar, Masoud Vedadi, Levon Halabelian

## Abstract

Cbl-b is a RING-type E3 ubiquitin ligase that is expressed in several immune cell lineages, where it negatively regulates the activity of immune cells. Cbl-b has specifically been identified as an attractive target for cancer immunotherapy due to its role in promoting an immunosuppressive tumor environment, and Nx-1607, is in phase I clinical trials for advanced solid tumor malignancies. Using a suite of biophysical and cellular assays, we confirmed potent binding of C7683 (an analogue of Nx-1607) to the full-length Cbl-b and its N-terminal fragment containing the TKBD-LHR-RING domains. To further elucidate its mechanism of inhibition, we determined the co-crystal structure of Cbl-b with C7683, revealing compound interaction with both the TKBD and LHR, but not the ring domain. Here, we provide structural insights into a novel mechanism of Cbl-b inhibition by a small-molecule inhibitor that locks the protein in an inactive conformation by acting as an intramolecular glue.

## INTRODUCTION

Post-translational modification by ubiquitination, the covalent conjugation of ubiquitin (Ub) to protein substrates, is an essential mechanism that regulates protein functions by altering their stability, cellular localization, activity, and protein-protein interactions in diverse biological processes. Ub modification requires consecutive actions of three classes of enzymes termed E1, E2 and E3^1,2^. Ubiquitin activating enzyme E1 activates and transfers Ub to a ubiquitin conjugating enzyme E2, resulting in covalent attachment of Ub to the E2 catalytic cysteine. Subsequently, an E3 ligase recruits both a protein substrate and E2-Ub thus facilitating Ub transfer from the E2 to the lysine residues of the protein substrate^3^. E3s thus confer substrate specificity during ubiquitination, and defects in their function are associated with many disease pathologies including cancers, metabolic disorders, and neurodegenerative diseases^4,5^.

The human Casitas B lymphoma-b (Cbl-b) protein is a member of the monomeric Cbl family of RING-type E3 ubiquitin ligases that are known to downregulate non-receptor and receptor protein tyrosine kinase (PTK) signaling by ubiquitinating and thereby targeting these kinases for proteasomal or lysosomal degradation^6,7^. The Cbl family of E3 ligases share a highly conserved N-terminal region that contains the structural elements required for ubiquitin ligase activity^8,9^, including a substrate recognition tyrosine kinase-binding domain (TKBD), a short linker helix region (LHR) and a RING finger domain (Fig. 1A). The TKB domain is composed of three subdomains, a 4-helix bundle (4H), a calcium binding domain with an EF-hand fold (EF hand), and a variant Src homology region 2 (SH2) domain (Fig. 1A), all of which form a unique phosphotyrosine-recognition module^10^ in which phosphorylated PTKs, such as ZAP-70 and Syk, are recruited for ubiquitination^11,12^. The RING finger domain directly associates with Ub conjugated E2 proteins and mediates target ubiquitination^13^.

**Figure 1:**
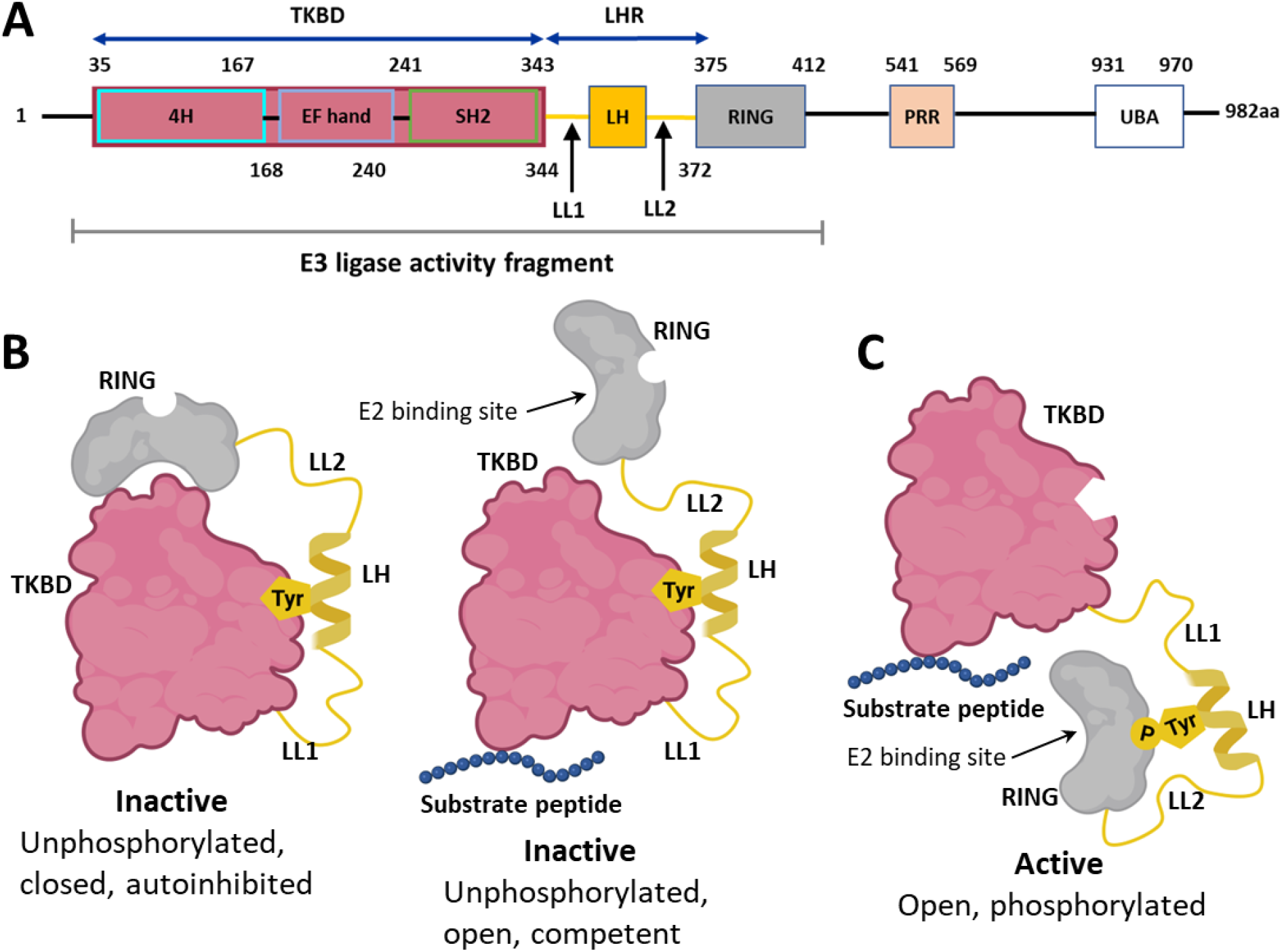
A model of the LHR-mediated regulation of Cbl proteins. **(A)** A schematic of the Cbl-b protein domains. The N-terminal fragment that confers the Cbl-b E3 ligase activity is indicated and contains 3 domains including: the TKBD colored in magenta (made up of three subdomains, 4H, EF hand and SH2), the LHR shown in yellow and the RING finger domain in grey. **(B)** A model of the unphosphorylated inactive Cbl-b N-terminal fragment of Cbl-b, colored and labeled according to the schematic in panel A. The LH is clamped onto the TKB domain, which restricts the movement of the RING finger domain to either closed and autoinhibited or open and competent of binding to E2. The conserved tyrosine (Tyr363 in Cbl-b) is labeled as Tyr. **(C)** A model of the phosphorylated active Cbl-b, colored and labeled according to panel A schematic. Upon Try363 phosphorylation, the LH is released from the TKBD, and the RING domain is flipped around 180 degrees and moved adjacent to the substrate. The phosphorylated tyrosine is labeled (PTyr). Figure adopted from Buetow et al. 2016^8^ and created with BioRender.com.

The LHR comprises an N-terminal loop (linker loop 1), a helix, and a C-terminal loop (linker loop 2) and contains a strictly conserved tyrosine residue (Tyr363 in Cbl-b) within the linker helix (LH) (Fig. 1) that is essential for regulating the Cbl proteins E3 ligase activity. Phosphorylation of this critical tyrosine activates the Cbl proteins, as its mutation to phenylalanine abolishes the E3 ligase activity^14^. In the unphosphorylated state, Cbl proteins are autoinhibited through the interaction of TKBD with the conserved tyrosine and several other residues within the linker helix, resulting in the RING finger domain being restricted to the side of the TKBD, opposite to the substrate binding site^8,15^ (Fig. 1B). In the inactive state, the RING finger domain can either be in a closed conformation unable to bind E2s, or in a substrate binding-induced open catalytically competent conformation that can interact with E2 proteins^8,13,16^ (Fig.1B). Upon phosphorylation, the LHR is released from the TKBD, which allows the protein to undergo conformational changes (Fig. 1C) that activate the protein. The activation process involves (i) unmasking of the RING finger domain from the TKBD, (ii) flipping of the RING domain around 180 degrees so it’s positioned adjacent to the substrate-binding site on TKBD, (iii) *p*Tyr-LHR forming interactions with the RING domain that stabilize E2-Ub during Ub transfer, and (iv) the phosphorylated tyrosine interacting directly with Ub thus generating a structural element adjacent to the RING finger domain that enhances its catalytic efficacy^3,15,16^.

In addition to the highly conserved N-terminus, Cbl-b and its closely related homologue c-Cbl contain a highly variable C-terminal extension that plays adaptor-like functions by mediating interactions with several protein families^8^. The C-terminus consists of a proline rich region (PRR) that mediates binding to SH3 domain-containing proteins, a tyrosine rich region that can recruit SH2 domain-containing proteins after phosphorylation of the tyrosines^17^, and an ubiquitin-associated domain (UBA) that is essential for homodimerization and heterodimerization between Cbl-b and c-Cbl^18^ and for binding ubiquitinated proteins, by Cbl-b only^18,19^.

Cbl-b is expressed in several immune cell lineages where it modulates both innate and adaptive immune responses, effectively playing a key role in the host defense mechanisms against pathogens and anti-tumor immunity^11^. Specifically, Cbl-b is a negative regulator of signaling pathways involving immune cell receptors, and it is particularly well-documented to play a central role in inhibiting effector T cell activation^20^. T cell activation requires both T cell receptor stimulation and CD28 receptor co-stimulation that potentiates Cbl-b ubiquitination and degradation. In the absence of CD28 co-stimulation, Cbl-b through ubiquitination and degradation-targeting of various signaling transducers (as reviewed by Tang et al. 2019^11^) inhibits T cell transcriptional activity hence promoting immune suppression in both adaptive and innate immunity^21^. This inhibitory effect is significant in cancer development and progression as it promotes an immunosuppressive tumor environment, making Cbl-b an attractive therapeutic target for cancer immunotherapy, as well as other human immune-related disorders including infections and autoimmune diseases^11,22^.

Using a high throughput screening, Sharp et al (2022) identified and developed a triazole based compound series targeting the E3 ligase activity of Cbl-b, including Nx-1607, which is currently in clinical trials as a cancer immunotherapeutic for the treatment of solid tumor malignancies (clinical trials.gov ID: NCT05107674)^23,24^. These compounds were shown to inhibit the interaction between the N-terminal E3 ligase fragment (TKBD-LHR-RING) of Cbl-b and UbcH5B-Ub (E2-Ub) complex in the presence of the Src kinase substrate (Patent number: WO2019148005), thus leading to the stimulation of immune cells in vitro and significant inhibition of tumors in vivo. However, the structural understanding of how these compounds interact with Cbl-b is lacking and hence their mechanism of action is not well characterized. Here, we report the co-crystal structure of the N-terminal E3 ligase fragment of Cbl-b in complex with a compound related to Nx-1607, which reveals that the Nx-1607 compound series bind to the TKBD-LHR unit of Cbl-b, locking the protein in an inactive conformation.

## RESULTS

### The TKB domain and the linker helix region are required for C7683 binding to Cbl-b

Sharp et al (2022) reported a series of triazole compounds targeting the E3 ligase activity of Cbl-b for cancer immunotherapy^23,24^. These compounds were shown to inhibit the interaction of Cbl-b with a ubiquitin-charged UbcH5B (E2-Ub) complex using a TR-FRET assay (Patent number: WO2019148005). In the absence of any structural information or other molecular mechanistic data, the mode of Cbl-b inhibition by these compounds is poorly understood. We therefore sought to understand the mechanism of action for a compound from this class, using C7683 as an exemplar, and in the process develop a toolkit for Cbl-b inhibitor characterization.

We first evaluated the binding activity of C7683, a representative triazole compound with isoindolin-1-one core structure that belongs to the same class as Nx-1607 reported in the patent application (WO2020264398), by differential scanning fluorimetry (DSF)^25^. Both full-length and the TKBD-LHR-RING (residues 38-427) proteins showed significant stabilization by C7683 at 3 μM concentration (Fig. 2 A-B), with ΔT_m_ values of 10 ± 0.4 and 12 ± 0.2 °C, respectively. The stabilization by C7683 was concentration dependent (Fig. 2 C-D), suggesting stoichiometric binding. This assay could be used to screen for new chemical series targeting the N-terminal domain of Cbl-b.

**Figure 2:**
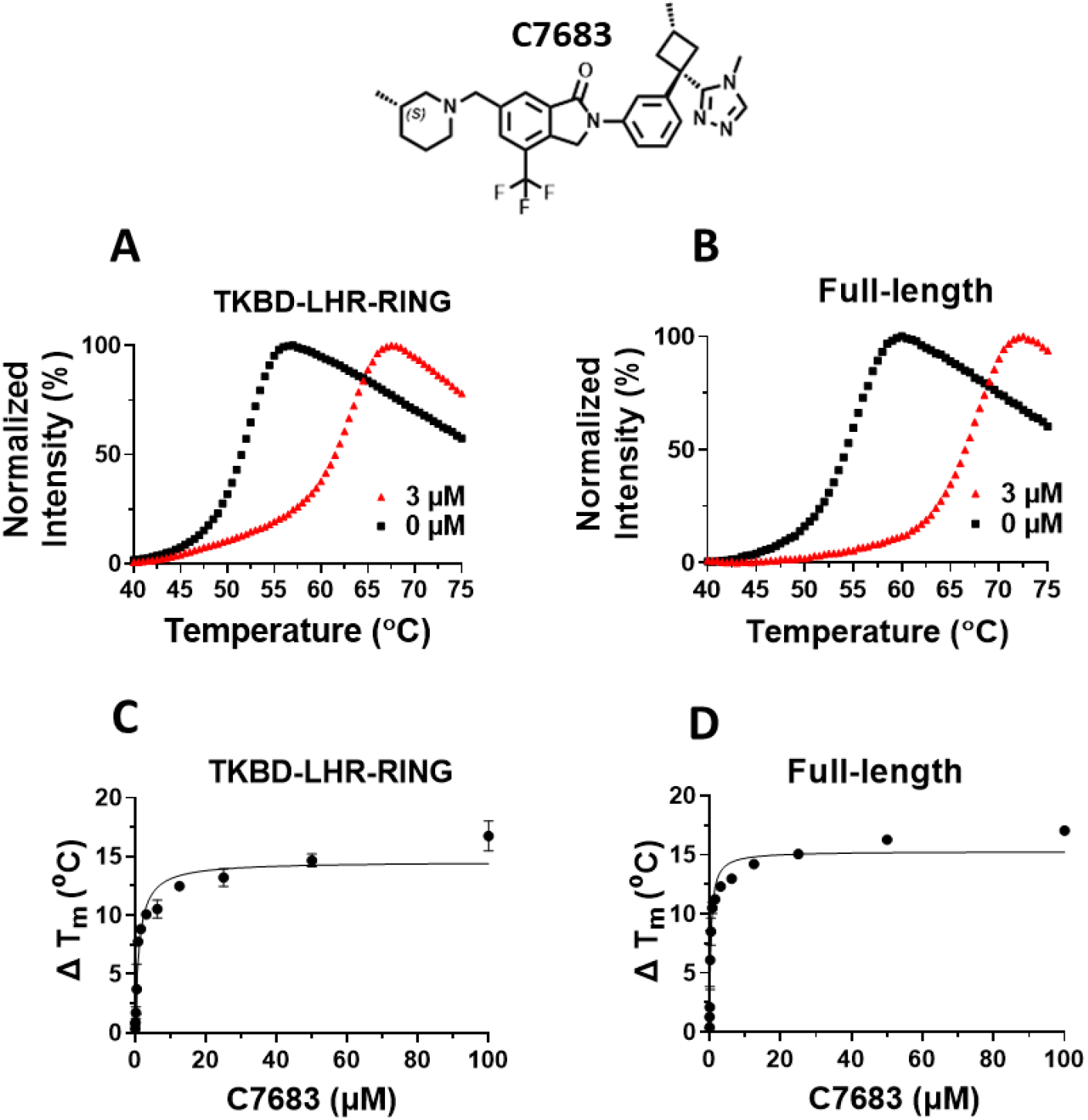
Assessment of C7683 binding to TKBD-LHR-RING fragment and full-length Cbl-b proteins by DSF. The chemical structure of C7683 is presented. C7683 stabilized (A) TKBD-LHR-RING and (B) full-length Cbl-b at 3 μM with ΔT_m_ values of 10 ± 0.4 and 12 ± 0.2 °C, respectively. The stabilization effect of C7683 was concentration dependent (C-D) for both constructs at concentrations ranging from 0.01 to 100 μM. All experiments were performed in triplicates (n=3).

Having confirmed the binding of C7683 to the full-length and the TKBD-LHR-RING Cbl-b proteins, we sought to identify the domain responsible for binding to C7683. As similar compounds were shown to inhibit the interaction of Cbl-b with UbcH5B-Ub complex in a TR-FRET based assay (Patent number, WO2019148005), we speculated that they may be binding to the RING domain of Cbl-b, which is an E2 binding module.

We first evaluated the binding of C7683 to the full-length, the TKBD-LHR-RING and the RING domain Cbl-b proteins using an orthogonal SPR assay. C7683 binds to both TKBD-LHR-RING and full-length proteins (Fig. 3 A-D), yielding K_D_ values of 8 ± 4 nM and 12 ± 6 nM respectively, but did not show any significant binding to the RING domain-only protein (Supp. Fig. S1A-B) suggesting that the TKBD/LHR regions are necessary for binding.

**Figure 3.**
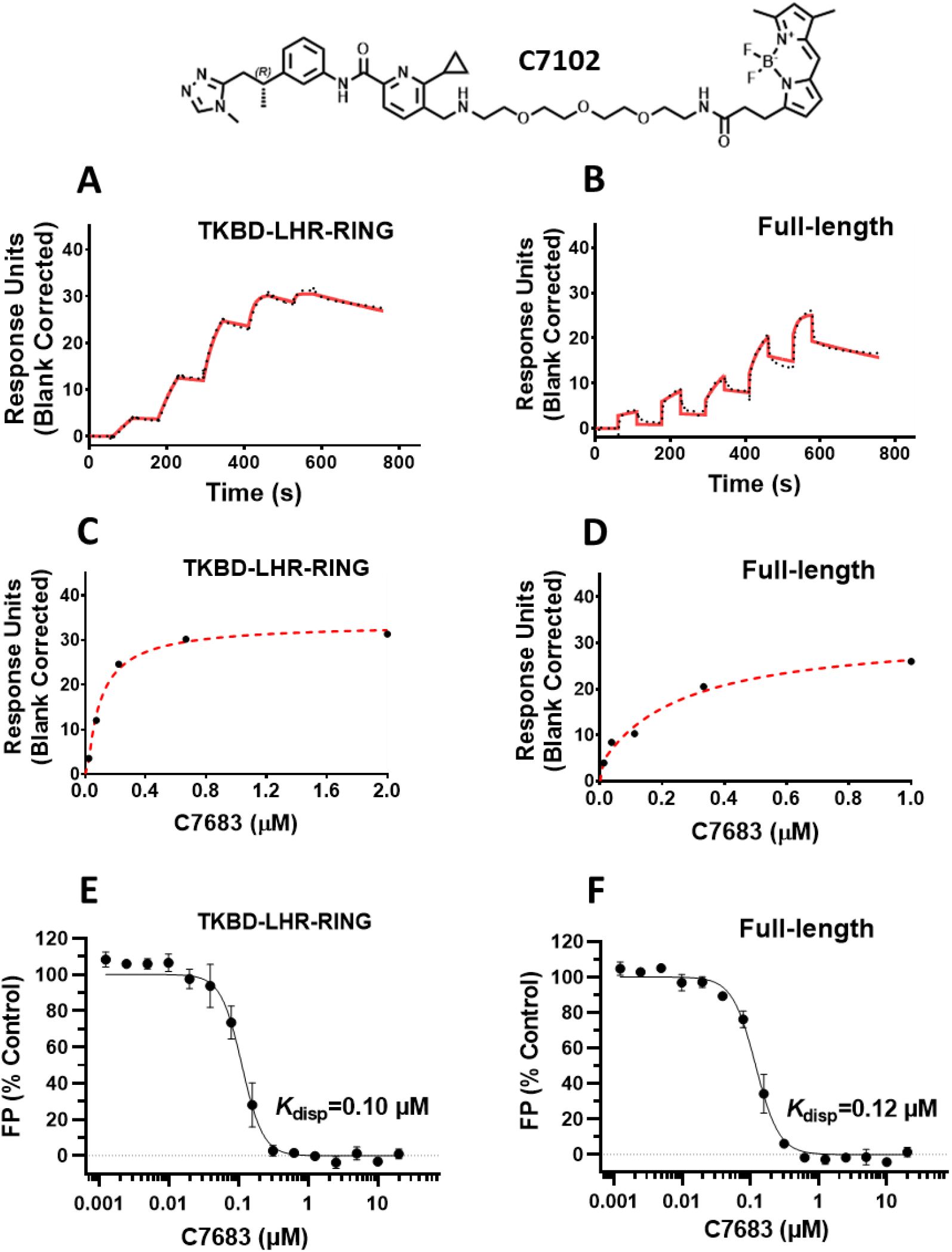
Assessment of C7683 binding to TKBD-LHR-RING and full-length Cbl-b proteins by SPR and a FP-based displacement assay. The chemical structure of the fluorescein-labeled probe (C7102) is presented. (A-D) Serially diluted C7683 was flowed over separately immobilized (A, C) TKBD-LHR-RING, and (B, D) full-length proteins. The representative sensorgrams for both sets of experiments are shown (solid red lines) with the kinetic fit (black dots), and the steady-state response (black circles) with the steady state 1:1 binding model fitting (red dashed line). C7683 was tested for competing with the fluorescein-labeled probe (C7102) for binding to (E) TKBD-LHR-RING and (F) full-length proteins at concentrations ranging from 0.001 to 20 μM. All experiments were performed in triplicate (n=3) and the calculated K_D_ and K_disp_ values are presented in supplementary Table 1.

To confirm this further, we selected a fluorescein-labeled compound (C7102, Fig. 3) related to C7683 and reported in a patent application (Compound 46, WO2020210508) to develop an FP-based displacement assay^26^. In this assay, C7683 displaced binding of C7102 to both TKBD-LHR-RING and full-length proteins in a concentration dependent manner (Fig. 3 E-F) but did not show any significant binding to the RING domain alone (Supp. Fig. S1C), confirming that the RING domain is not involved in C7683 binding. The DSF ΔT_m_, the SPR K_D_ and the FP K_*disp*_ values are shown in Supplementary Table S1.

### C7683 stabilizes Cbl-b protein in the cells

As C7683 has shown a large thermal shift in DSF assay, we wanted to determine if C7683 binds Cbl-b protein in cells and show similar thermal stabilization. We developed a HiBIT target engagement cellular thermal shift assay (CETSA), which measures shifts in protein thermal stability following compound treatment. In this assay, HiBIT-tagged full length Cbl-b was expressed in HEK293T cells and cells were treated with increasing C7683 concentrations, exposed to a temperature gradient, and lysed in a buffer containing LgBIT protein. HiBIT complementation with LgBIT results in formation of the functional NanoLUC luciferase enzyme, which produces bioluminescence signal following substrate addition, and is dependent on the amount of soluble HiBIT protein at each temperature. C7683 showed significant dose-dependent stabilization of Cbl-b protein in cells (Fig. 4). Significant thermal stabilization was observed at concentration as low as 1 nM, with thermal shifts of > 15 °C observed at 1 μM to 10 μM compound concentration, consistent with our biophysical thermal shifts assay results (Fig. 2).

**Figure 4.**
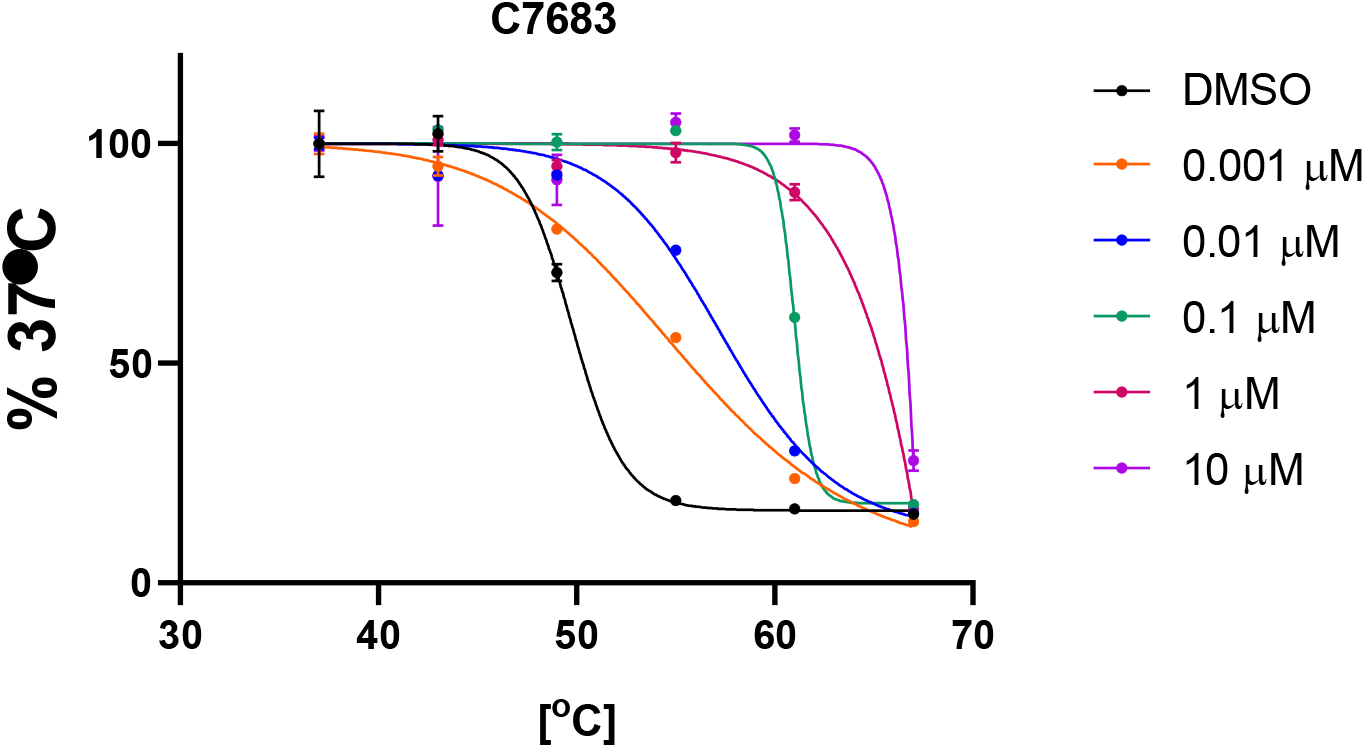
C7683 potently stabilizes Cbl-b in cells - HiBIT CETSA assay. As described in the material and methods, the amount of unaggregated HiBIT-tagged Cbl-b was quantified. C7683 significantly stabilized the Flag-Hb-Cbl-b protein in a dose-dependent manner compared to DMSO control. The results are MEAN+/-SD, n=4.

### C7683 binds at the interface of the TKBD and LHR of Cbl-b

To further characterize the C7683 interaction with Cbl-b and understand its mechanism of action, we co-crystallized the N-terminal E3 ligase fragment of Cbl-b containing the TKBD, LHR and RING domains (residue range 38 to 427) in complex with C7683. Initial co-crystallization trials with a TKBD-LHR-RING protein containing a Tyr360 mutation to Phe (similar to PDB ID: 3ZNI^3^) generated poorly diffracting crystals in our hands. To improve the crystal packing and obtain better diffracting crystals, we introduced three-point mutations on the surface of the Cbl-b TKB domain by mutating residues K51, K55, R325 to alanines (the mutant protein referred to here as 3mCbl-b), which yielded crystals diffracting to 1.8 Å resolution. These mutations are located far from the substrate binding site as well as the regulatory regions involved in Cbl-b activation. The co-crystal structure of 3mCbl-b-C7683 (PDB: 8GCY) was determined in P2_1_2_1_2_1_ space group symmetry, with one copy of the Cbl-b in the crystal asymmetric unit bound to C7683. Supplementary Table S2 summarizes the crystallographic data collection, refinement, and validation statistics.

A well-resolved electron density map was observed for the entire C7683 molecule (Supp. Fig. S2) and Cbl-b protein, except for residues 339 to 353 in the flexible linker helix region (LHR). C7683 was nestled in an interface between the 3 TKBD subdomains and the LHR (Fig. 5A-B), consistent with our biophysical studies, which showed that the compound binding does not involve the RING domain. C7683 binds in an extended conformation, making extensive interactions with the residues of all three TKBD subdomains, the LHR and coordinating a nearby water molecule (HOH10) (Fig. 5C-D). The methyl-substituted piperidine ring is nested in a pocket between the 4H and the SH2 subdomains, where it makes hydrophobic and van der Waals contacts with Cys289, Thr265, Pro71 and Pro72, in addition to a hydrogen bonding interaction between its nitrogen atom (N31) and the side chain oxygen (OE1) of Glu268 in the SH2 domain. The Isoindolin-1-one ring also located between 4H and SH2 subdomains, making hydrophobic interactions with atoms from Thr144, Lys145, L148 in the 4H domain and Leu264 in the SH2, and hydrogen bonds to the backbone nitrogen of Phe263 in the SH2 domain through its oxygen atom. The phenyl ring is nestled between the side chains of Leu222 and Met366 in the SH2 domain and the LHR respectively, while the methyl-substituted cyclobutane ring makes hydrophobic contacts with the side chains of Thr219, and Leu287 in the Ca^2+^ binding EF hand and the SH2 domains, respectively. The triazole ring interacts primarily with the SH2 domain and the LHR; it makes two hydrogen bonds to the sidechain oxygen of Tyr260 and a nearby water molecule (HOH10). The triazole ring is also involved in π-π stacking interaction with the side chain ring of the critical Tyr363 in the LHR, which plays a crucial role in regulating Cbl-b’s ligase activity, as well as hydrophobic interactions with Ser218 and Gly367 (Fig. 5C-D).

**Figure 5.**
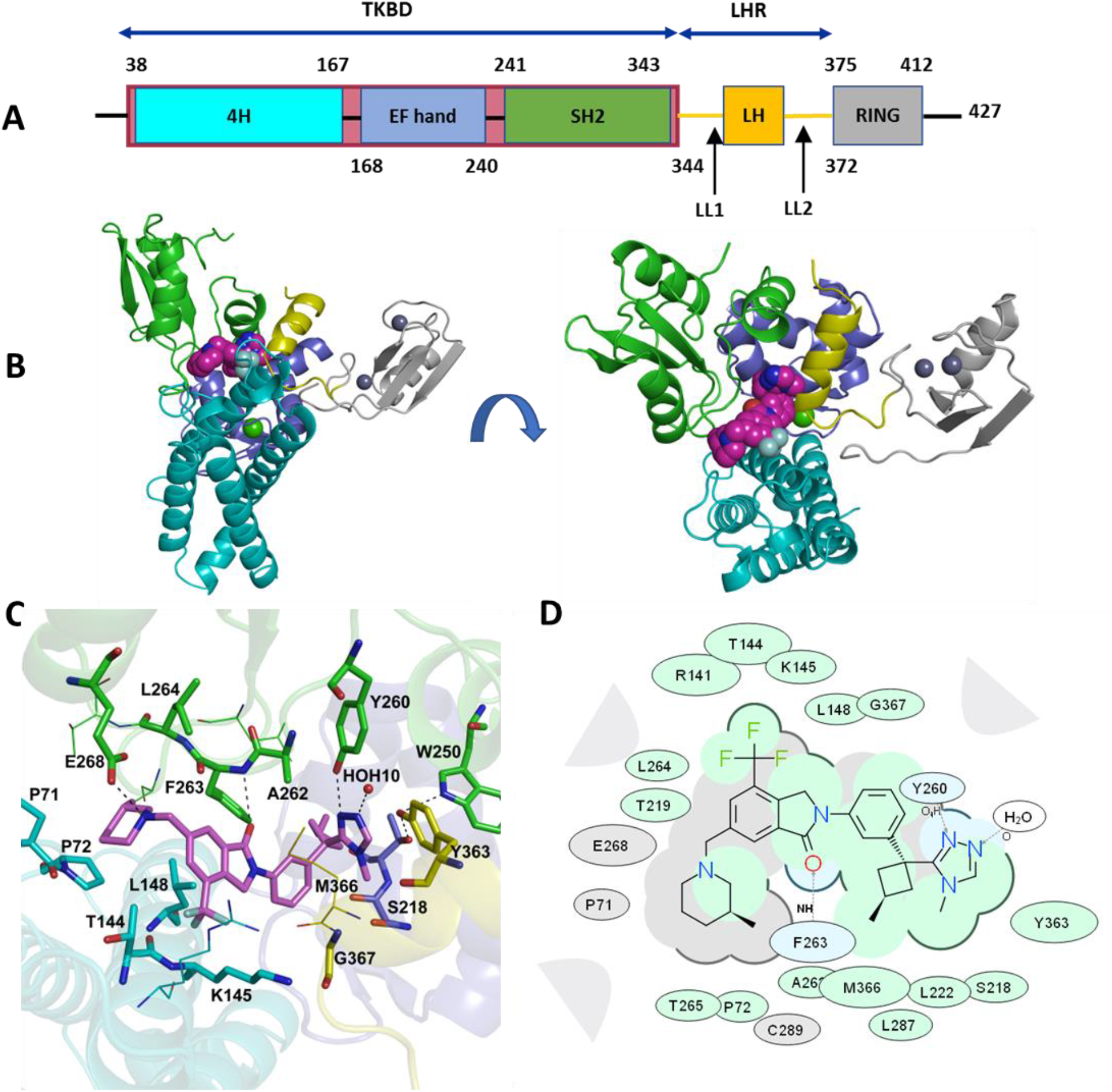
X-ray crystal structure of Cbl-b N-terminal fragment bound to C7683. **(A)** Domain organization of the crystallized Cbl-b N-terminal fragment. **(B)** Side and top views of the 3mCbl-b-C7683 showing the compound nestled in an interface between the three TKBD subdomains and the linker helix region (LHR) of Cbl-b. The TKBD subdomains are shown as cartoon representation and colored according to the domain organization in panel **A**, and the compound is shown as magenta spheres. **(C)** A close-up view of the C7683 binding site. C7683 forms a network of interactions with a water molecule (red sphere), the three TKBD subdomains and the LHR. Hydrogen bond interactions are shown in black dashes. The compound is shown as magenta sticks and key interacting residues are rendered as sticks. **(D)** A 2D depiction of the protein-compound interactions. Blue shading represents hydrogen bonding residues and interactions, green shading hydrophobic interactions and grey shading other van der Waals contacts. (Figure generated with ICM-Pro (v. 3.8-2c, MolSoft CA, USA)).

### C7683 locks Cbl-b in an inactive conformation

The co-crystal structure of 3mCbl-b-C7683 revealed that the protein adopts an inactive conformation having the LHR docked between the TKBD subdomains and the RING domain adopting a conformation that limits its binding to the E2 ligase^3,8,15,16^. Figure 6 compares the 3mCbl-b-C7683 structure with those of inactive and active conformations of Cbl-b and other Cbl proteins within the family. The RING finger domain adopts different conformations (orientation and position) in all the five structures compared. The 3mCbl-b-C7683 structure has a unique RING domain orientation compared to other Cbl protein forms (Fig. 6B and Supp. Fig. 3A). Compared to the unbound c-Cbl structure (PDB ID: 2Y1M)^16^, the 3mCbl-b-C7683 RING domain appears to be in a partially open orientation, with some of the RING domain E2-interacting residues (Ile375, Trp399, Phe418) that are known to interact with the TKBD in the closed Cbl conformation being fully or partially exposed to the solvent (Supp. Fig. S3 B-C). However, based on its orientation and proximity to TKBD/LHR, the 3mCbl-b-C7683 RING domain would not be capable of binding to an E2 ligase, and therefore it can be considered to be in a closed inactive state.

**Figure 6.**
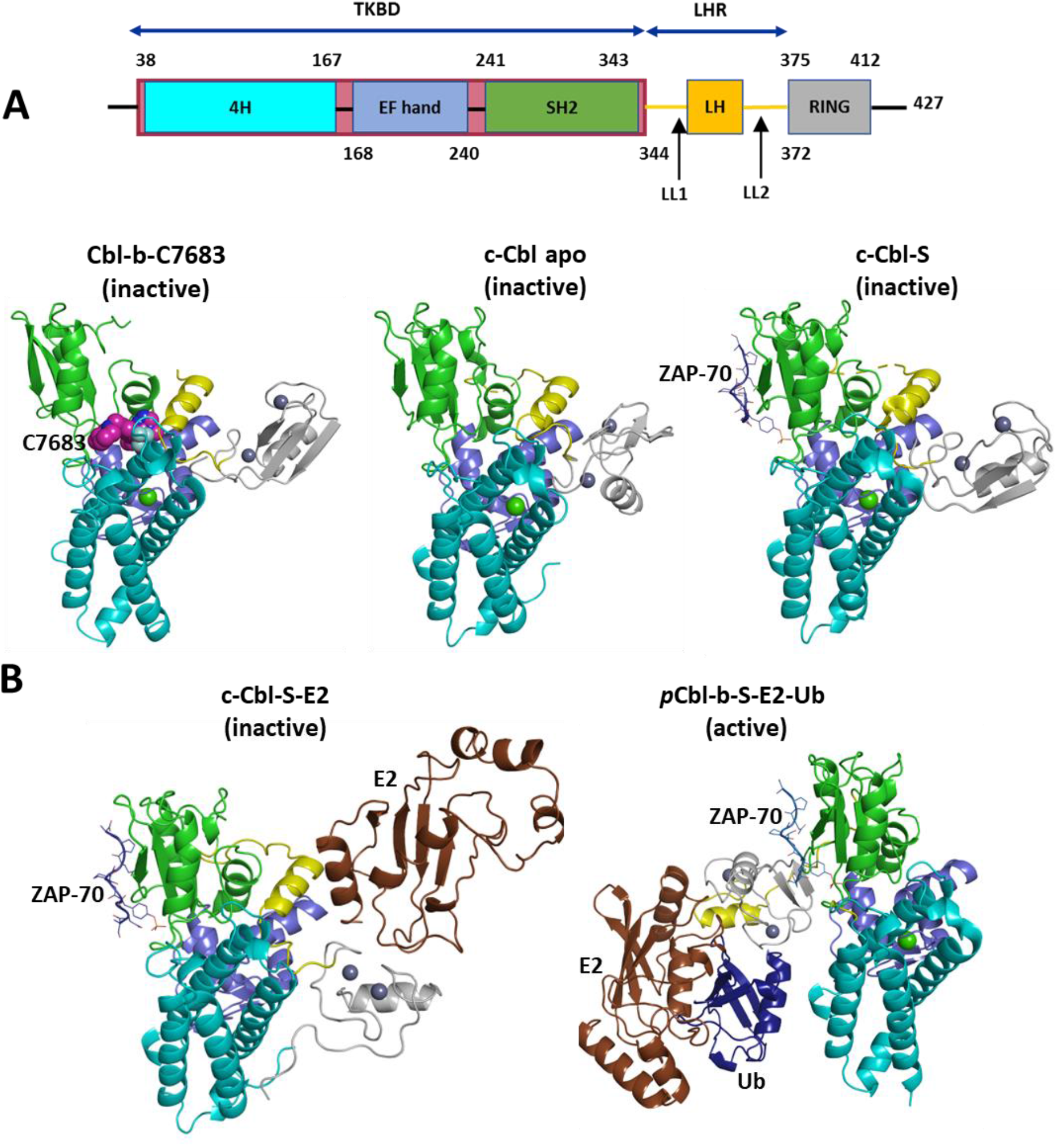
Structures of Cbl protein family switching between an active (Tyr363 phosphorylated) and inactive (Tyr363 non-phosphorylated) conformations. **(A)** Domain organization of the N-terminal fragment of Cbl-b represented in the structures. **(B)** Comparison of Cbl-b-7683 structure with (i), the c-Cbl apo structure (PDB ID: 2Y1M)^16^, (ii), the c-Cbl in complex with a ZAP-70 substrate peptide (c-Cbl-S) (PDB ID: 2Y1N)^16^, (iii) the c-Cbl in complex with ZAP-70 peptide and UbcH7 E2 ligase (c-Cbl-S-E2) (PDB ID: 1FBV)^13^, and (iv) the *p*Tyr363 Cbl-l in complex with ZAP-70 peptide, UbcH5B E2 ligase, and ubiquitin (*p*Cbl-b-S-E2-Ub) (PDB ID: 3ZNI)^3^. The CBL domains are colored according to the domain schematic in panel A, the C7683 compound is shown as magenta spheres, ZAP-70 peptide is rendered as a blue loop and sticks, E2 ligases are depicted as brown cartoons and ubiquitin is rendered as deep blue cartoons.

As described above, C7683 interacts with the LHR through residues including Met366 and the conserved Try363 (Fig. 5B). The activation of Cbl-b is dependent on the phosphorylation of Try363, which releases the LHR from interaction with the TBKD, allowing the protein to undergo large conformational changes that promote effective ligase activity (Fig. 6B & Supp. Fig. S4). These changes include exposure of the E2-binding surface of the RING domain to facilitate efficient interaction with the E2 ligase, and bringing the E2-bound RING domain into close proximity to the TKBD substrate binding region of Cbl-b^8,16^ (Fig. 1). The compound binding at the interface between the TKBD and LHR, as well as its specific interaction with the Tyr363 in the flexible LHR, locks the protein in an inactive conformation that prevents phosphorylation of Tyr363, which is crucial for Cbl-b activation.

A comparison of the compound-bound 3mCbl-b-C7683 structure with that of unbound inactive Cbl-b (PDB ID: 3VGO)^15^ shows that the TKBD/LHR interface is not altered upon compound binding (Supp. Fig. S4A). In substrate-bound and activated forms of Cbl proteins (PDB ID: 2Y1N, PDB ID: 1FBV and PDB ID: 3ZNI)^3,13,16^, the SH2 domain loop containing Phe263 is displaced by approximately 4 Å following both substrate binding and tyrosine phosphorylation^16^ (Supp. Fig. S4B). C7683 makes numerous contacts with this loop and fills the gap created in the TKBD domain of inactive Cbl-b conformation that otherwise would be occupied by SH2 domain loop following substrate binding and Tyr363 phosphorylation (Supp. Fig. S4). Thus, C7683 would not only prevent Tyr363 phosphylation for Cbl-b activation, but it would also block the conformational changes resulting from substrate binding to the TKBD domain of Cbl-b.

## Discussion

In this study, we have structurally and biochemically characterized the mechanism of Cbl-b inhibition by C7683, a new class of E3 ligase inhibitor related to Nx-1607, which is currently in clinical trials for treatment of advanced solid tumor malignancies. We show that C7683 binds Cbl-b with high affinity and significantly stabilizes the protein in both *in vitro* and in cellular assays. Importantly, we show that C7683 interacts with the TKBD and LHR of Cbl-b, thereby acting as an intramolecular glue by locking the protein in an inactive (closed) conformation.

This presents a novel mechanism for the inhibition of ubiquitination activity of an E3 ligase by locking the non-phosphorylated form of the protein in a closed conformation, and without directly binding to the E2-binding RING domain. Other E3 ligases, mainly the HECT domain E3 ligases such as WWP1, WWP2, NEDD4^27–29^ also adopt similar closed autoinhibited state, in which its multiple WW domains sequester HECT using a multi-lock mechanism and phosphorylation of linker or cancer associated mutation relieve autoinhibition. As compounds targeting E2-binding HECT domain or substrate binding domains of these type of E3 ligases are limited, similar to this novel mechanism, HECT type E3 ligases could be inhibited by compounds stabilizing the inactive closed conformation.

In the case of Cbl-b, C7683 induced a large thermal shift as observed *in vitro* by DSF assay (Fig. 2) and in cells by CETSA assay (Fig. 4). These simple thermal shift assays could be used to identify novel inhibitors of Cbl-b as well as for other E3 ligases such as HECT E3 ligases acting via the same mechanism.

Here, we have developed target enabling resources for the human Cbl-b protein, an attractive target for cancer immunotherapy and other human immune disorders. Specifically, we have generated structural data and developed biophysical and cellular assays to support future drug development efforts to selectively inhibit Cbl-b and other Cbl E3 ligases involved in human diseases.

## METHODS

### Biophysical binding characterization of C7683

#### Protein expression and purification

Biophysical experiments were carried out using three Cbl-b protein variants: the full-length (1-982aa), the N-terminal E3 ligase fragment consisting of TKBD-LHR-RING (38-427aa) and the RING finger domain (351-426aa). The full-length Cbl-b gene was cloned into an in-house Baculovirus expression vector, pFBD-BirA, a derivative of pFastBac Dual vector from Invitrogen, containing an N-terminal AviTag, a C-terminal hexa-His tag and co-expression of BirA. Biotinylated Cbl-b protein was expressed in *Sf9* insect cells following a protocol described by Hutchinson and Seitova^30^ and using biotin-supplemented media. The N-terminal E3 ligase fragment and the RING domain genes were cloned into a pET28-derived in-house *E. coli* expression vector, p28BIOH-LIC, containing an N-terminal AviTag and a C-terminal hexa-His tag. The two proteins were co-expressed with BirA contained in a pBirAcm vector (Avidity LLC) in *E. coli* BL21 cells using biotin-supplemented media. All the proteins were purified using Nickel affinity chromatography, followed by size exclusion chromatography using a HiLoad™ 26/60 Superdex™ 75 gel filtration column (GE Healthcare).

#### Compound synthesis

The C7683 and fluorescein-labeled C7102 compounds were synthesized as reported in patent applications (WO2020210508, WO2020264398).

#### Thermal shift assay by Differential Scanning Fluorimetry (DSF)

DSF^25^ experiments were performed using a Roche LightCycler 480 II in sealed Axygen 384-well plates (PCR-384-LC480). Reactions (20 μL) contained 0.1 mg/mL TKBD-LHR-RING and full-length Cbl-b, in 30 mM Hepes pH 7.5, 150 mM NaCl, 0.5mM TCEP, and SYPRO Orange (5000× SYPRO Orange stock in DMSO, Invitrogen). C7683 was tested at a serially diluted range from 0.01 to 100 μM with a final concentration of 2% DMSO. Thermal denaturation was monitored from 20 to 95 °C at a rate of 4 °C increase per minute, and data points were collected at 0.5 °C intervals.

#### Analysis of binding by Surface plasmon resonance (SPR)

*In vitro* binding analyses by SPR were carried out using a Biacore T200 (GE Health Sciences Inc.) instrument at 20 °C. 20 mM Hepes, pH 7.5, 100 mM NaCl, 0.5 mM TCEP, 0.01% Triton X-100 was used for all the experiments. Biotinylated TKBD-LHR-RING, full-length, and RING domain Cbl-b proteins were immobilized yielding approximately 4000, 9000, and 2000 response units (RU), respectively, on a flow cell of a Streptavidin-coated (SA) sensor chip (GE healthcare) according to manufacturer’s directions while another flow cell was left blank for reference subtraction. The C7683 compound was diluted in the same buffer to yield a 2% final DMSO solution and was serially diluted 3-fold. The compound was then flowed over the sensor chip using a single cycle Kinetics mode at 50 μL/min. Contact time (i.e., association phase) was 50 s and disassociation time was 180 s. Buffer with 2% DMSO was used for blank injections; and buffers containing 1 to 3% DMSO were used for solvent correction injections. Binding constants were acquired from the double referenced (i.e., reference subtraction and blank injection subtraction) single cycle data using Biacore T200 Evaluation Software 3.1.

#### Fluorescence polarization (FP)-based displacement assay

The FP-based displacement experiments^26^ were performed in 384-well polypropylene small volume black microplates (Cat. # 784209, Greiner) in a total volume of 20 μL per well at room temperature (i.e., 23 °C). C7683 compound was added to the reaction mixture containing 850 nM TKBD-LHR-RING, full-length and RING domain Cbl-b proteins, and 40 nM of the fluorescein-labelled probe (C7102) in buffer (20 mM Hepes, pH 7.4, 150 mM NaCl, 0.01% Triton X-100, 0.5 mM TCEP), and were incubated for 30 min at room temperature. Final DMSO concentration was 2%. The resulting FP signals were measured using a BioTek Synergy 4 (BioTek, Winooski, VT) at the excitation and emission wavelengths of 485 nm and 528 nm, respectively. The obtained FP values were blank subtracted and are shown as the percentage of control. All the experiments were carried out in triplicate (n=3) and the presented values are the average of replicates ± standard deviation. FP data were visualized using GraphPad Prism software 8.0 (La Jolla, CA).

### Cellular Target Engagement

We performed a Cbl-b cellular thermal shift assay (CETSA) using NanoLuc split luciferase technology (Promega). HEK293T cells were plated in 6-well plates (4^e5^ cells/ml) and 4 h later transfected with 0.2 μg of N-terminally HiBIT–tagged full-length Cbl-b and 1.8 μg of empty plasmid using X-tremeGene XP transfection reagent, following manufacturer’s instructions. Next day cells were trypsinized and resuspended in OptiMEM (no phenol red) at a density of 2e5/ml. After compound or DMSO addition (the same DMSO concentration in each sample) cells were transferred to 96-well pcr plates (50 μl/well), incubated for 1 h at 37 °C and heated at indicated temperatures in thermocycler for 3 min. After 3 min at RT, 50 μl of LgBIT solution (200 nM LgBIT, 2% NP-40, prot. inhibitors in OptiMEM no phenol red) was added and incubated at RT for 10 min. Next, 25 μl of NanoGlo substrate (8 μl/ml) was added, mixed gently and 20 μl was transferred to 384 white plates in quadruplicates and bioluminescence signal was read.

### Protein Crystallography

#### Cbl-b gene cloning, protein expression and purification

The N-terminal TKBD-LHR-RING fragment (residues 38-427) of human Cbl-b gene (UniProtKB Q13191) was cloned into an in-house pET28-derived *E. coli* expression vector called pET28-MHL, yielding a construct with an N-terminal His_6_-tag followed by a TEV cleavage site. Crystal engineering was then carried out to reduce the surface entropy by mutating residues K51, K55 and R325 into alanines to promote crystallization. The triple mutant Cbl-b fragment 3mCbl-b protein was expressed overnight at 16 °C in *E. coli* BL21 (DE3) pRARE2 cells.

Protein purification was performed by immobilized Nickel ion affinity chromatography, followed by size exclusion chromatography using a HiLoad™ 16/60 Superdex™ 75 gel filtration column (GE Healthcare). The protein was then subjected to TEV protease digestion overnight at 4 °C to remove polyhistidine purification tag. The protein was further purified by anion exchange chromatography using a SOURCE™ 15 Q column (MILLIPORE SIGMA) and eluted in the final protein buffer containing 20mM Tris-HCl pH 7.5, 165 mM NaCl, 2.5% Glycerol and 2mM TCEP. Protein fractions containing pure Cbl-b protein as confirmed by SDS-PAGE were pooled and concentrated using 10 kDa cutoff spin columns (Millipore), and the final protein concentration determined using the NanoDrop UV-Vis spectrophotometer (Thermo Scientific), with the protein extinction coefficient of 62340 M-1cm-1 computed from the amino acid sequence using Expasy ProtParam (https://web.expasy.org/protparam/).

#### Co-crystallization of Cbl-b in complex with C7683

To generate Cbl-b-C7683 co-crystals, purified 3mCbl-b protein at 10.7 mg/ml (0.237 mM) concentration was mixed with 3 times molar excess of C7683 (0.712 mM) and incubated at room temperature for 15 minutes prior to setting crystallization trays. Crystallization was carried out using the sitting drop vapor-diffusion method by mixing the protein-C7683 complex with an equal volume of the reservoir solution over 100 μL reservoir. Crystals were observed within 72 hours at 18 °C in a precipitant solution containing 0.1 M Imidazole, 0.1 M MES monohydrate pH 6.5, 0.09 M Sodium nitrate, 0.09 M Sodium phosphate dibasic, 0.09M Ammonium sulfate, 12.5% v/v MPD; 12.5% PEG1000; 12.5% w/v PEG 3350.

#### Diffraction data collection, structure determination and refinement

Crystals in the crystallization mother liquor that already contained cryoprotectant (MPD) were directly cryo-cooled in liquid nitrogen. Diffraction data were collected on beamline 24-ID-E at the Advanced Photon Source in Argonne National Laboratory. The diffraction data were processed with HKL3000^31^ and structure was solved by molecular replacement in Phaser^32^ using the apo Cbl-b crystal structure (PDB ID: 3VGO)^15^ as a starting model. The model was refined by alternating cycles of manual rebuilding in Coot^33^ and refinement with Refmac^34^ within the CCP4 crystallography suite^35^. The structure was validated using the Molprobity server^36^, and the molecular graphics images were rendered using PyMOL^37^. The model coordinates are deposited in the RCSB PDB under PDB ID: 8GCY.

## ACCESSION NUMBERS

Atomic coordinates and structure factors for the reported crystal structures have been deposited in the Protein Data bank under the accession number: 8GCY

## ACKNOWLEDGEMENTS

We would like to thank Peter Loppnau, Almagul Seitova, Ashley Hutchinson for insect cell protein expression, and Albina Bolotokova for compound dissolution and management. This work is based upon research conducted at the Northeastern Collaborative Access Team beamlines, which are funded by the National Institute of General Medical Sciences from the National Institutes of Health (P30 GM124165). This research used resources of the Advanced Photon Source, a U.S. Department of Energy (DOE) Office of Science User Facility operated for the DOE Office of Science by Argonne National Laboratory under Contract No. DE-AC02-06CH11357. Structural Genomics Consortium is a registered charity (no: 1097737) that receives funds from Bayer AG, Boehringer Ingelheim, Bristol Myers Squibb, Genentech, Genome Canada through Ontario Genomics Institute [OGI-196], EU/EFPIA/OICR/McGill/KTH/Diamond Innovative Medicines Initiative 2 Joint Undertaking [EUbOPEN grant 875510], Janssen, Merck KGaA (aka EMD in Canada and US), Pfizer and Takeda.

## REFERENCES

1. Varshavsky, A. The ubiquitin system, an immense realm. Annu. Rev. Biochem. 81, 167–176 (2012).

2. Hershko, A. & Ciechanover, A. The ubiquitin system for protein degradation. Annu. Rev. Biochem. 61, 761–807 (1992).

3. Dou, H., Buetow, L., Sibbet, G. J., Cameron, K. & Huang, D. T. Essentiality of a non-RING element in priming donor ubiquitin for catalysis by a monomeric E3. Nat. Struct. Mol. Biol. 20, 982–986 (2013).

4. Petroski, M. D. The ubiquitin system, disease, and drug discovery. BMC Biochem. 9 Suppl 1, S7 (2008).

5. Hoeller, D. & Dikic, I. Targeting the ubiquitin system in cancer therapy. Nature 458, 438–444 (2009).

6. Mohapatra, B. et al. Protein tyrosine kinase regulation by ubiquitination: critical roles of Cbl-family ubiquitin ligases. Biochim. Biophys. Acta 1833, 122–139 (2013).

7. Swaminathan, G. & Tsygankov, A. Y. The Cbl family proteins: ring leaders in regulation of cell signaling. J. Cell. Physiol. 209, 21–43 (2006).

8. Buetow, L. et al. Casitas B-lineage lymphoma linker helix mutations found in myeloproliferative neoplasms affect conformation. BMC Biol. 14, 76 (2016).

9. Thien, C. B. F. & Langdon, W. Y. c-Cbl and Cbl-b ubiquitin ligases: substrate diversity and the negative regulation of signalling responses. Biochem. J. 391, 153–166 (2005).

10. Meng, W., Sawasdikosol, S., Burakoff, S. J. & Eck, M. J. Structure of the amino-terminal domain of Cbl complexed to its binding site on ZAP-70 kinase. Nature 398, 84–90 (1999).

11. Tang, R., Langdon, W. Y. & Zhang, J. Regulation of immune responses by E3 ubiquitin ligase Cbl-b. Cell. Immunol. 340, 103878 (2019).

12. Lupher, M. L., Songyang, Z., Shoelson, S. E., Cantley, L. C. & Band, H. The Cbl phosphotyrosine-binding domain selects a D(N/D)XpY motif and binds to the Tyr292 negative regulatory phosphorylation site of ZAP-70. J. Biol. Chem. 272, 33140–33144 (1997).

13. Zheng, N., Wang, P., Jeffrey, P. D. & Pavletich, N. P. Structure of a c-Cbl-UbcH7 complex: RING domain function in ubiquitin-protein ligases. Cell 102, 533–539 (2000).

14. Levkowitz, G. et al. Ubiquitin ligase activity and tyrosine phosphorylation underlie suppression of growth factor signaling by c-Cbl/Sli-1. Mol. Cell 4, 1029–1040 (1999).

15. Kobashigawa, Y. et al. Autoinhibition and phosphorylation-induced activation mechanisms of human cancer and autoimmune disease-related E3 protein Cbl-b. Proc. Natl. Acad. Sci. U. S. A. 108, 20579–20584 (2011).

16. Dou, H. et al. Structural basis for autoinhibition and phosphorylation-dependent activation of c-Cbl. Nat. Struct. Mol. Biol. 19, 184–192 (2012).

17. Schmidt, M. H. H. & Dikic, I. The Cbl interactome and its functions. Nat. Rev. Mol. Cell Biol. 6, 907–918 (2005).

18. Peschard, P. et al. Structural basis for ubiquitin-mediated dimerization and activation of the ubiquitin protein ligase Cbl-b. Mol. Cell 27, 474–485 (2007).

19. Kozlov, G. et al. Structural basis for UBA-mediated dimerization of c-Cbl ubiquitin ligase. J. Biol. Chem. 282, 27547–27555 (2007).

20. Lutz-Nicoladoni, C., Wolf, D. & Sopper, S. Modulation of Immune Cell Functions by the E3 Ligase Cbl-b. Front. Oncol. 5, 58 (2015).

21. Liu, Q., Zhou, H., Langdon, W. Y. & Zhang, J. E3 ubiquitin ligase Cbl-b in innate and adaptive immunity. Cell Cycle Georget. Tex 13, 1875–1884 (2014).

22. Augustin, R. C., Bao, R. & Luke, J. J. Targeting Cbl-b in cancer immunotherapy. J. Immunother. Cancer 11, e006007 (2023).

23. Sharp, A. et al. A first-in-human phase 1 trial of nx-1607, a first-in-class oral CBL-B inhibitor, in patients with advanced solid tumor malignancies. J. Clin. Oncol. 40, TPS2691–TPS2691 (2022).

24. Discovery and optimization of Cbl-b inhibitors (https://www.nurixtx.com/wp-content/uploads/2022/09/Nurix-CBL-B-DOT-Talk_JK.pdf).

25. Niesen, F. H., Berglund, H. & Vedadi, M. The use of differential scanning fluorimetry to detect ligand interactions that promote protein stability. Nat. Protoc. 2, 2212–2221 (2007).

26. Allali-Hassani, A. et al. Fluorescence-based methods for screening writers and readers of histone methyl marks. J. Biomol. Screen. 17, 71–84 (2012).

27. Verdecia, M. A. et al. Conformational flexibility underlies ubiquitin ligation mediated by the WWP1 HECT domain E3 ligase. Mol. Cell 11, 249–259 (2003).

28. Chen, Z. et al. A Tunable Brake for HECT Ubiquitin Ligases. Mol. Cell 66, 345-357.e6 (2017).

29. Wang, Z. et al. A multi-lock inhibitory mechanism for fine-tuning enzyme activities of the HECT family E3 ligases. Nat. Commun. 10, 3162 (2019).

30. Hutchinson, A. & Seitova, A. Production of Recombinant PRMT Proteins using the Baculovirus Expression Vector System. J. Vis. Exp. JoVE (2021) doi:10.3791/62510.

31. Minor, W., Cymborowski, M., Otwinowski, Z. & Chruszcz, M. HKL-3000: the integration of data reduction and structure solution--from diffraction images to an initial model in minutes. Acta Crystallogr. D Biol. Crystallogr. 62, 859–866 (2006).

32. McCoy, A. J. et al. Phaser crystallographic software. J. Appl. Crystallogr. 40, 658–674 (2007).

33. Emsley, P. & Cowtan, K. Coot: model-building tools for molecular graphics. Acta Crystallogr. D Biol. Crystallogr. 60, 2126–2132 (2004).

34. Murshudov, G. N., Vagin, A. A. & Dodson, E. J. Refinement of macromolecular structures by the maximum-likelihood method. Acta Crystallogr. D Biol. Crystallogr. 53, 240–255 (1997).

35. Winn, M. D. et al. Overview of the CCP4 suite and current developments. Acta Crystallogr. D Biol. Crystallogr. 67, 235–242 (2011).

36. Chen, V. B. et al. MolProbity: all-atom structure validation for macromolecular crystallography. Acta Crystallogr. D Biol. Crystallogr. 66, 12–21 (2010).

37. DeLano, W. & Schrödinger, L. PyMOL. Retrieved Http://www.pymolorgpymol.

