## Supplementary Figures for "Probing the mechanism of Cbl-b inhibition by a small-molecule inhibitor"

**Vijayaratnam Santhakumar**

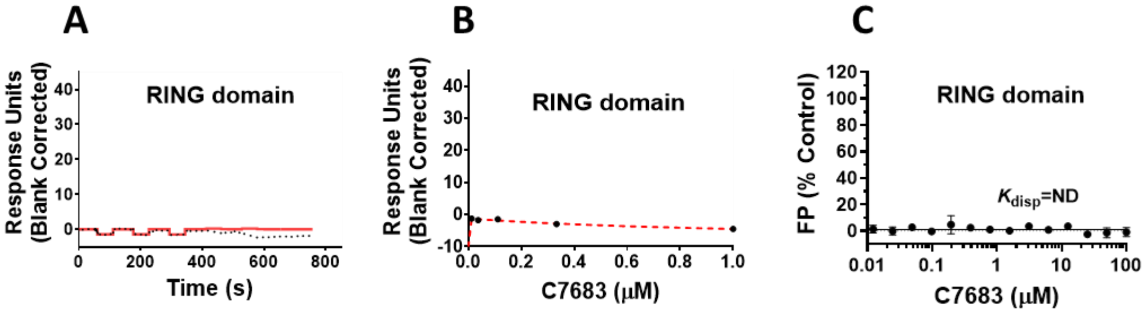


**Figure S1. Assessment of C7683 binding to Cbl-b RING domain by SPR and FP probe displacement. (A-B)** Serially diluted C7683 was flown over immobilized RING domain protein (351-426aa). Representative sensorgram is shown in (solid red lines) with the kinetic fit (black dots) in (A), and the steady-state response (black circles) with the steady state 1:1 binding model fitting (red dashed line) in (B). **(C)** C7683 was tested for competing with the fluorescein-labeled probe (C7102) for binding to Cbl-b RING domain. C7683 didn't show any binding to the RING domain by SPR and FP. All experiments were performed in triplicates (n=3).

**Table S1. Summary of binding and peptide displacement assays.** All values are the average ± standard deviation from experiments presented in figures 2 and 3 above (n=3). SPR $K_D$ values are from kinetic fitting.

| **Target** | **Displacement** $K_{disp}$ **(μM)** | **DSF** $\Delta T_m$ **(°C)** | **SPR** $K_D$ **(nM)** |
|---|---|---|---|
| TKBD-LHR-RING | 0.10 ± 0.02 | 10 ± 0.4 | 8 ± 4 |
| Full-length | 0.12 ± 0.02 | 12 ± 0.2 | 12 ± 6 |

**Table S2: Data collection and refinement statistics**

| | **3mCbl-b-C7683** |
|---|---|
| PDB ID | 8GCY |
| Wavelength (nm) | 0.9791 |
| Resolution range (Å) | 43.68 - 1.81 (1.84 - 1.81)[a] |
| Space group | P 21 21 21 |
| Unit cell (Å)<br>(˚) | 47.937, 76.093, 105.464<br>90.00, 90.00, 90.00 |
| Total reflections | 388627 |
| Unique reflections | 35624 |
| Multiplicity | 10.9 (9.9) |
| Completeness (%) | 98.6 (98.3) |
| Mean I/sigma(I) | 36.00 (1.92) |
| R-merge | 0.041 (0.785) |
| R-meas | 0.067 (0.990) |
| R-pim | 0.020 (0.306) |
| CC1/2 | 0.997 (0.827) |
| CC* | 0.999 (0.951) |
| Reflections used in refinement | 33780 |
| Reflections used for R-free | 1795 |
| R-work | 0.197 |
| R-free | 0.220 |
| CC (work) | 0.961 |
| CC (free) | 0.943 |
| Number of non-hydrogen atoms | |
| Macromolecule | 3039 |
| Ligands/ions | 54 |
| Solvent | 143 |
| Protein residues | |
| RMS (bonds) | 0.0094 |
| RMS (angles) | 1.4512 |
| Average B-factor | |
| Macromolecule | 36.249 |
| Ligands/ions | 29.018 |
| Solvent | 36.461 |

[a]Values in the parentheses are for the highest-resolution shell.

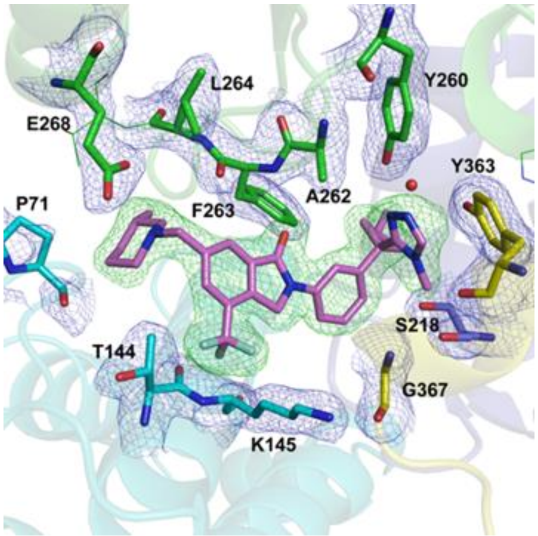


**Figure S2. Electron density map of C7683 bound to the TKBD subdomains and LHR of Cbl-b.** The protein residues in the binding site are shown as sticks and are colored based on their respective domains as portrayed in figure 5A. A coordinating water molecule is rendered as a red sphere and the measured 2Fo-Fc electron density map around some of the highlighted residues in the vicinity of the compound is shown as blue mesh, contoured at 1.0σ level. C7683 electron density omit map (Fo-Fc) is shown as green mesh contoured at 3σ level and the C7683 compound is rendered as magenta sticks.

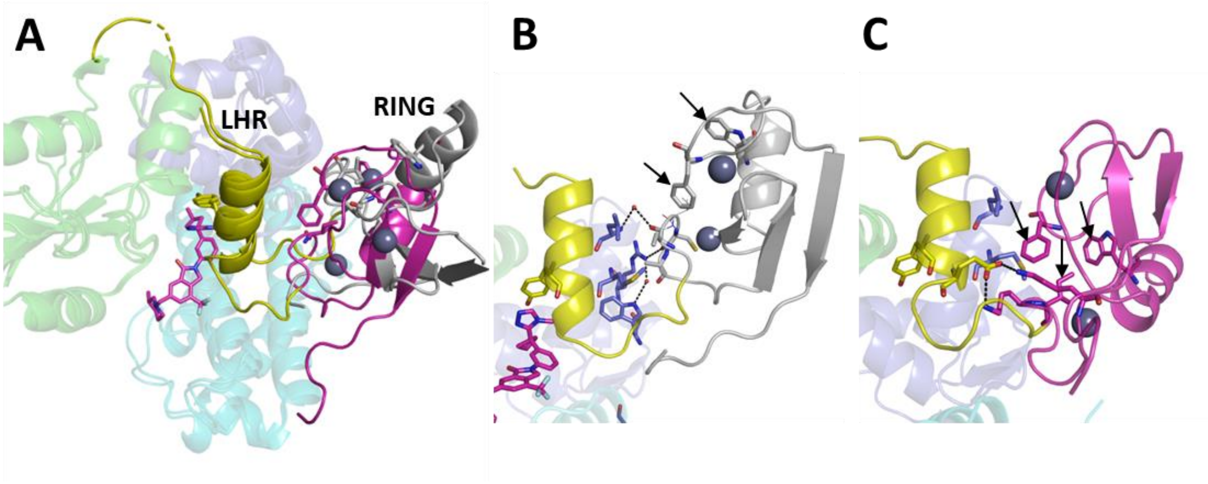


**Figure S3. The unique RING domain conformation in the 3mCbl-b-C7683 structure. (A)** A comparison of the RING finger domain configuration in the 3mCbl-b-C7683 (grey) compared to c-Cbl apo (magenta, PDB ID: 2Y1M)[16] structures. The RING domain in the C7683-bound structure adopts a unique conformation. **(B)** The interactions of the RING domain with TKBD/LHR in the 3mCbl-b-C7683 structure. Unique hydrogen bonding interactions involving the RING domain and TKBD are observed, and some of the RING domain E2-interacting residues (Trp399, Phe418, see black arrows) that are known to interact with the TKBD/LHR in the autoinhibited closed Cbl conformation are exposed. **(C)** The interactions of the RING finger domain with the TKBD/LHR in the closed c-Cbl apo structure (PDB ID: 2Y1M)[16]. Three of the E3-interacting RING domain residues (Ile383, Trp408, Phe418, see black arrows) that interact with TKBD/LHR in the closed Cbl conformation form hydrophobic interactions with Leu219 and Met222 of TKBD and LHR domains respectively[16].

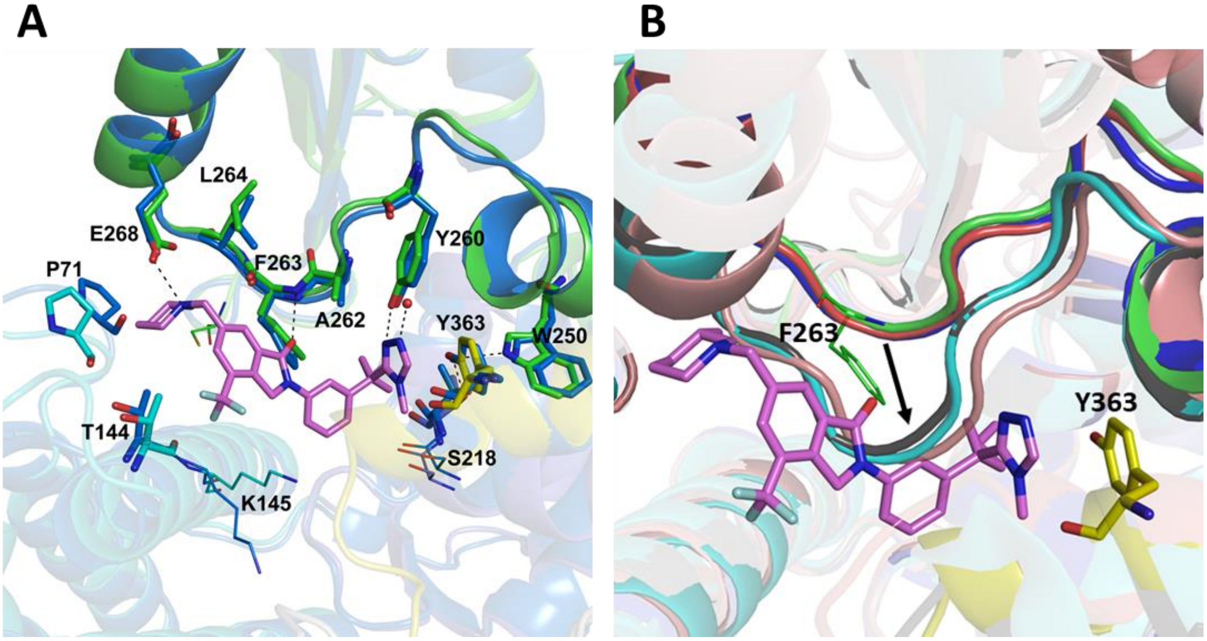


**Figure S4. Comparison of the compound binding pocket of 3mCbl-b-C7683 with that of active and inactive Cbl structures. (A)** An overlay of the 3mCbl-b-C7683 (green, C7683 magenta sticks) and the apo Cbl-b (blue, PDB ID: 3VGO)[15] structures (both inactive). The Phe263-bearing loop, as well as the amino acid sidechains in the vicinity of the compound overlap well in both structures, indicating that binding of the compound does not induce changes in the binding site. **(B)** A superposition of C7683-bound, active and inactive Cbl structures. The C7683-bound structure (green), Cbl-b apo structure (blue, PDB ID: 3VGO)[15] and c-Cbl apo structure (red, PDB ID: 2Y1M)[16] have the Phe263-bearing loop in the same conformation, which represents an inactive closed Cbl conformation. The c-Cbl protein in complex with a ZAP-70 substrate peptide structure (cyan, PDB ID: 2Y1N)[16], the c-Cbl protein in complex with ZAP-70 peptide and UbcH7 E2 ligase structure (grey, PDB ID: 1FBV)[13], and the *p*Tyr363 Cbl-l structure in complex with ZAP-70 peptide, UbcH5B E2 ligase, and ubiquitin (brown, PDB ID: 3ZNI)[3] have the Phe263-bearing loop that is crucial for compound binding shifted by ~4 Å as indicated by the black arrow - which represents an open, inactive substrate-bound or active (phosphorylated) conformation.
